# Theory on the power-law distribution of the Michaelis-Menten enzyme kinetic parameters

**DOI:** 10.1101/2023.09.16.558070

**Authors:** Rajamanickam Murugan

## Abstract

We show on the biophysical basis that the observed enzyme kinetic parameters related to the substrate binding, product turnover and catalytic efficiency follow power-law type density functions. These finding are validated with the available datasets on various enzymes across different substrates. The product turnover rates and catalytic efficiencies seems to follow a bimodal type density functions in line with the single molecule experiments on enzyme catalysis which can be explained by a sum of two different power-law type density functions. The curve geometry of the density functions is decided by the underlying biophysical factors and the location of the peak is dictated by the natural selection pressure.

## INTRODUCTION

Enzymes are core biological catalysts which drive various metabolic pathways essential for the existence of life on earth [1-3]. The catalytic mechanism of enzymes can be modelled by the Michaelis-Menten scheme (**Fig 1**) in which the substrate reversibly binds the enzyme to form the enzyme-substrate complex that reduces the activation energy barrier and drives the substrate towards its product. The catalytic power and efficiency of an enzyme can be characterized by various rates associated with the binding-unbinding of the substrate and subsequent catalysis and product release.

**FIGURE. 1.**
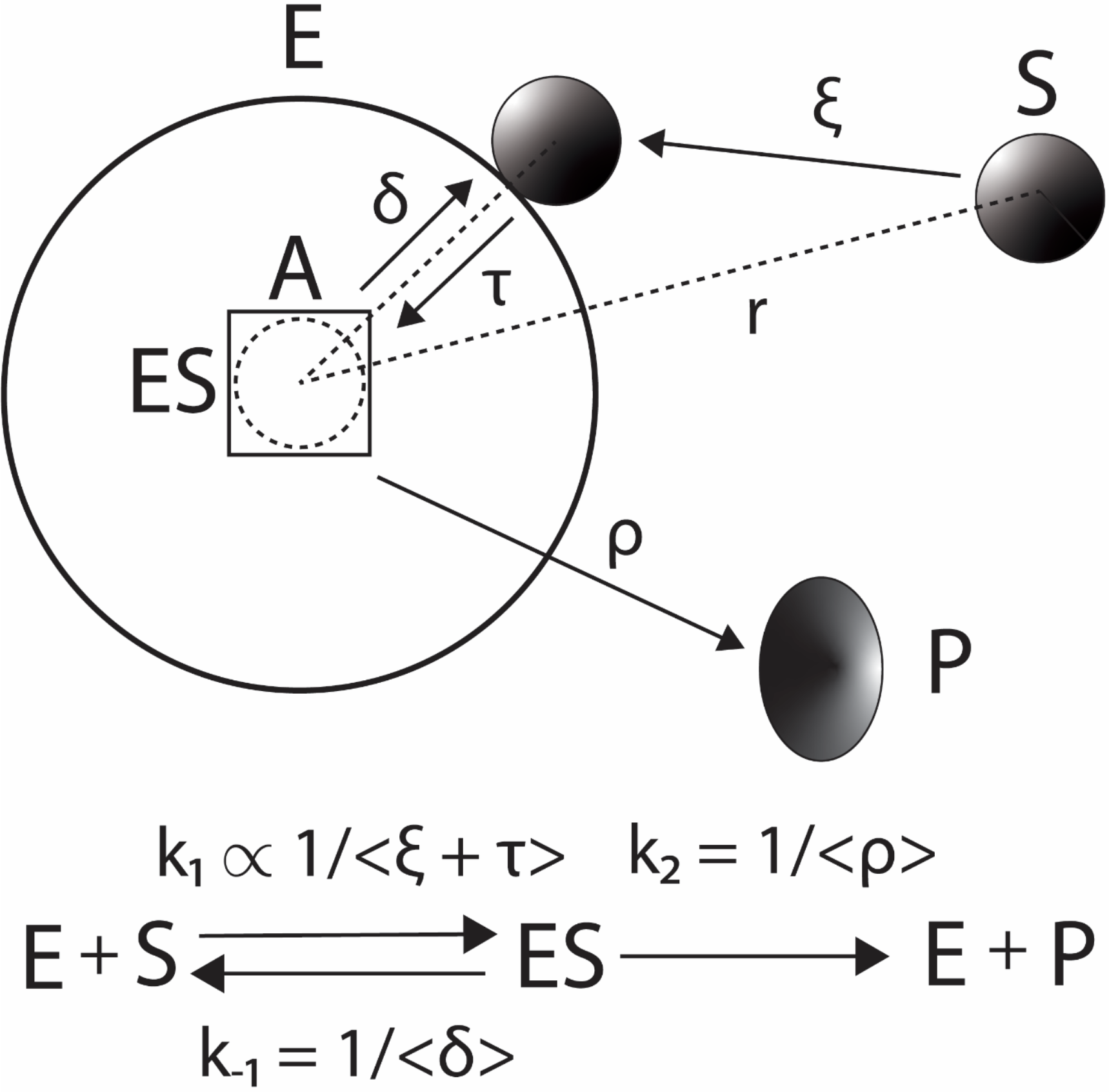
Various time components in the Michaelis-Menten enzyme kinetic scheme. The substrate S first reach the active site (A) of enzyme (E) via three-dimensional diffusion in time ξ and then binds there via several conformational changes and bonding interactions in time τ. As a result, the overall bimolecular collision rate associated with the formation of enzyme substrate complex (ES) will be *k*_1_ ∝ 1/ ⟨ *ξ* + *τ*⟩. Further, ρ and δ are the time components related to the product (P) formation and substrate dissociation respectively.

The rate constants connected with the binding (k_1_, 1/mols/lit/second), unbinding (k_-1_, 1/second) of substrate and the product release (k_2_, 1/second) can be summarized by the parameters viz. Michaelis-Menten constant (K_M_ = (k_-l_ + k_2_) / k_1_, measured in mols/lit), the catalytic turnover rate (k_cat_ = k_2_) and the catalytic efficiency k_E_ = k_2_/K_M_ (1/mols/lit/second) [3]. It is an intriguing question how nature designs and optimizes these parameters so that enzymes work at maximum efficiency and speed well within the metabolic framework. Availability of datasets on these kinetic parameters across various enzymes over different organisms, experimental conditions and substrates allows us to computationally address this question in detail [4-8]. However, the underlying physics behind the origin and shape of the observed probability distributions of these parameters is sill not clear. In this context, we will show that the enzyme parameters K_M_, k_cat_ and K_E_ follow approximately power-law type density functions which are fine-tuned by both underlying biophysical factors apart from the generally proposed natural selection pressure [5]. We will further validate our theoretical findings with the available data on these enzyme kinetic parameters.

Let us define the random variables associated with K_M_, k_cat_ and K_E_ as (*γ, u, v*) respectively such that K_M_ = < ϒ >, k_cat_ = <u> and K_E_ = <v> are their ensemble averages. We further define the random time variables (*σ, ρ, δ*) such that their ensemble averages are <δ> = 1 / k_-l_, <ρ> = 1/k_2_ and <σ> = 1/c k_1_. Here c is the scaling factor related to the concentration component of the bimolecular collision rate k_l_ so that σ is measured in time units, δ is the dissociation time and ρ is the product formation and release time. With these, one can define *γ* related to K_M_ in terms of the random times associated with the binding-unbinding and catalytic turnover as *γ = c*(*σ* (*ρ + δ*)*/ ρ δ*) (**Fig 1**). Further, one finds that u = 1/ρ and v = u/ ϒ. Here the realizations of (*σ*, ρ. *δ*) as well as (*γ, u, v*) depend on several core factors viz. hydrodynamic size of the enzyme, conformational fluctuations of the active site, hydrodynamic size and conformation of the substrate, bonding and non-bonding interactions of the substrate at the active site, and the presence of inhibitors apart from the physical factors such as temperature, pH and ionic strength of the reaction medium. Here the core factors are generally optimized along the evolutionary process.

### THEORETICAL METHODS

The reaction dynamics can be modelled as the diffusion over a potential. Let us consider a linear lattice of length L = |X_R_ - X_A_| with boundaries at (X_R_, X_A_) such that x = (X_R_ < X_A_) which represents the reaction coordinate. Here the absorbing boundary is at x = X_A_ and the reflecting boundary is at x = X_R_. The probability density function associated with a random walker (here it is the reaction dynamics) under potential *f*(x) confined inside such lattice obeys the following Fokker-Planck equation [9-11].

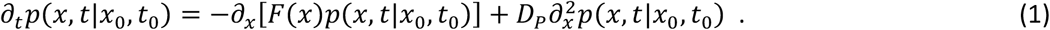

Here *F*(*x*) *= − df* (*x*)*/dx* is the force generated by the potential *f*(x), *p*(*x, t1x*_0_, *t*_0_) is the probability of finding a random walker at (x, t) with the condition that the random walker was at x = x_0_at t = t_O_, and D_P_ is the one-dimensional phenomenological diffusion coefficient associated with the reaction dynamics. The PDE given **Eq (1)** is a separable one with the substitution *p*(*x, t*) *= H*(*x*)*exp*(*− λ t − f*(*x*)*/D*_*p*_) leading to the canonical form [12].

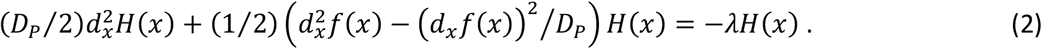

Since the differential operator in the left hand side of **Eq (2)** is a Hermitian one, the solution to **Eq (1)** can be expanded in terms of biorthogonal set of eigenfunctions [9-11] *Ψ*_*m*_(*x*) *= H*(*x*)*exp*(*− f*(*x*)*/D*_*p*_) corresponding to the given boundary conditions as follows.

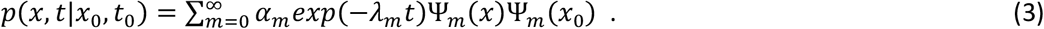

The absorbing boundary at x = X_A_ is set by *Ψ* _*m*_(*X*_*A*_) *=* 0 and the reflecting boundary at x = X_R_ is set by 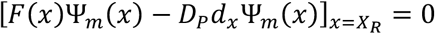. Here *λ* _*m*_ is the mr^th^ eigen value corresponding to the mr^th^ eigenfunction *Ψ* _*m*_(*x*) and α _*m*_ is the respective normalization constant. Clearly, *p*(*x, ∞ 1x*_0_, *t*_0_) *=* 0 since the random walker escape out of (X_R_, X_A_) over time with a probability of one. The distribution of the escape or first passage times from (X_R_, X_A_) starting from X_R_ ≤ x_O_ ≤ X_A_ can be derived from **Eq (3)** as 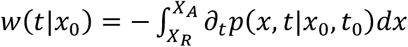 [10, 11]. Since the first eigenvalue *λ*_*1*_ will be the leading one, we obtain the asymptotic approximation *w*(*t1x*_0_)*∝exp*(*−λ*_*1*_*t*)*Ψ*_*1*_(*x*_0_). In general, the distribution of the escape times approximately follows an exponential type as *w*(*t*) *≅ λ*_*1*_*exp*(*−λ*_*1*_*t*) especially for large timescales.

In this background, we first consider the time required for the binding of substrate with the enzyme σ which is a sum of at least two different time components viz. the time required for the three-dimensional diffusion mediated collision of the substrate with enzyme *ξ* and the time required for further structural rearrangements and specific bond formation at the active site *τ* so that *σ = ξ + τ* (**Fig. 1**). From the theory of diffusion-controlled reactions [13, 14], one can obtain the probability *p*(*r, ξ*) of finding the substrate molecules at a distance *r* from the centre of static enzyme sphere at time *ξ* as follows.

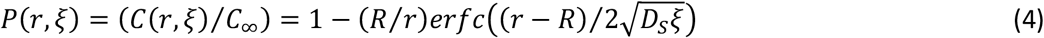

Here 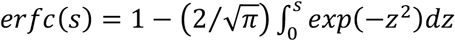 [15] is the complementary error function, R is the reaction radius which is the sum of the radii of gyration of the substrate and enzyme, D_S_ is the translational diffusion coefficient of the substrate, *C*_*∞*_ is the bulk concentration of the substrate and *C*(*r, ξ*) is the concentration of the substrate at a distance *r* from the enzyme at time *ξ*. Using **Eq (4)**, one can obtain the distribution of *ξ* as follows.

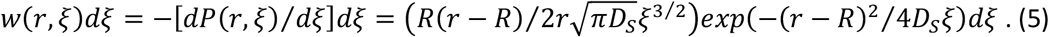

From **Eq (5)**, one can deduce the scaling *w*(*r, ξ*) *∝ ξ* ^*−3/2*^ as the leading one. The subsequent binding, dissociation and product formation steps are all generally orchestrated via structural rearrangements, bonding and non-bonding interactions between the substrate and the amino acid side chains of the enzyme active-site. Therefore, one can model the binding, unbinding and product formation steps as one-dimensional diffusion of the system over a double well potential with finite activation energy barrier along the respective reaction coordinate within the framework of Kramer’s theory [16]. In line with **Eqs (1-3)**, one can approximate the distributions of the first passage time components i.e., (*τ, ρ, δ*) in the definition of (*γ, u, v*) with appropriate boundary conditions as follows.

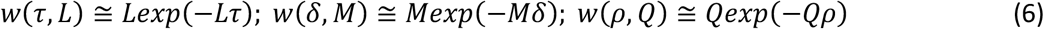

Here L, M and Q are the respective parameter exponents. Since the distribution function associated with *ξ* decays much faster than τ, one can approximate as σ ≅ *τ*. This means that the overall binding time *σ* approximately follows an exponential type density function. Noting that *γ ≅* (*τ* (*ρ + δ*)*/ ρ δ*) and *δ « ρ* since the product formation is generally the rate limiting step [1, 3], one obtains *γ ≅* (*τ / δ*). Using the transformation rule *δ d γ = d τ* one can derive the distribution of *γ* as follows.

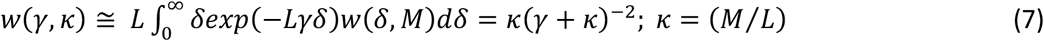

**Eq (7)** suggests that K_M_ i.e., γ values obtained by repeated *in vitro* experiments under various conditions will follow a power law type density function with diverging mean and variance. Remarkably, when *γ → ∞*, then such repeated realizations of *γ* will exhibit a tail as *w*(*γ, κ*) *∝ y* ^*−2*^. Further, the transformation *y = ln*(*γ*) leads to *exp*(*y*)*dy = d γ* and one finds the following density function for *y*.

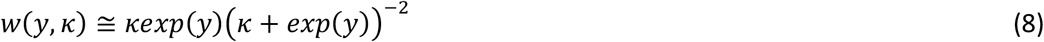

This density function has the mean value 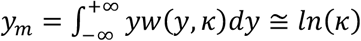 and the variance 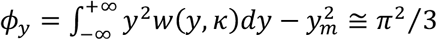 that is remarkably independent on *κ*. This is the most probable value of the sample variance (denoted as var(y)) of y. When *y → ∞*, then one finds *w*(*y, κ*) *∝ exp*(*− y*) that is the signature of a power law trend in the log-transformed space.

Consider the definition of inhibitor constant K_l_ = k_i_ / k_-i_. Here k_i_ is the bimolecular collision rate (1/mol/lit/second) between the enzyme and the inhibitor molecule and k_-i_ is the dissociation rate of the enzyme-inhibitor complex. In line with our arguments on K_M_, we define *K*_*I*_ = ⟨ *η* ⟩ where *η = τ*_*i*_ */δ*_*i*_ is the random variable associated with K_l_, *τ* _*i*_ is the time required to for the binding of inhibitor at the enzyme active site and *δ*_*i*_ is the time required for the dissociation of the enzyme-inhibitor complex. Since both *τ*_*i*_ and *δ*_*i*_ follow exponential type distributions, one can conclude that η, values obey a power law distribution similar to ϒ given in **Eqs (7-8)**. When *η → ∞*, then *w*(*η, κ*) *∝ η* ^*−2*^. Upon defining q = ln(η), one can show that *w*(*q, κ*) *∝ exp*(*− q*) as *q → ∞*. Since both the substrate and inhibitor share the common active site of the enzyme, one can also predict a positive correlation between K_M_ and K_l_ i.e., *γ* and *η*. However, unlike the direct experimental estimation of K_M_ values from the double reciprocal plots [17], K_l_ values will be estimated indirectly [1, 3] by first estimating the K_M_ values at different inhibitor concentrations [I] and then K_l_ will be extracted [3] from the quasi steady state relationship *K*_*M,I*_ *= K*_*M*_(*1 +* [*I*]*/K*_*I*_). Here K_M_ corresponds to the zero-inhibitor concentration [I] = 0. Since *ln*(*K*_*I*_) *= ln*(*K*_*M*_) *− ln*(*K*_*M,I*_) *+ ln*(*K*_*I*_ *+* [*I*]) and, y and q share a common density function, the most probable value of the experimentally observed variance of q will be approximately three times of *ϕ*_*y*_ as *ϕ*_*q*_ *≅ π*^*2*^. This result can be obtained by applying the variance operator on the right hand side of *ln*(*K*_*I*_).

Now we consider k_cat_ that is defined as *k*_cat_ = *k*_2_ = <*u*> = 1/<*p*>. From the definition of K_M_ = (k_−1_+ k_2_) / k_1_, one finds that *ln*(*k*_*2*_) *= ln*(*k*_*1*_) *+ ln*(*K*_*M−*_ *k*_*−1*_*/k*_*1*_). Since *ln*(*k*_*1*_) *» ln*(*K*_*M−*_*k*_*−1*_*/k*_*1*_) and the term (*K*_*M*_ *−k*_*−1*_*/k*_*1*_) *= k*_*2*_*/k*_*1*_ is similar to the definition of *ϒ*, the density function corresponding to z = ln(u) will be similar to **Eq (8)** as follows.

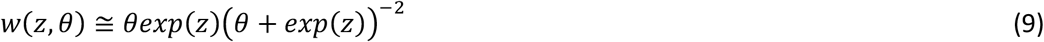

**Eq (9)** suggests that when *z → ∞* then *w*(*z, θ*) *∝ exp*(*−z*) with mean *z*_*m*_ *≅ ln*(*θ*) and variance *ϕ*_*z*_ *> ϕ*_*y*_. Noting the facts that *k*_E_ = *k*_2_ / K_M_ = <*v*> where v = u / ϒ, and *ln*(*k*_*E*_) *= ln*(*k*_*2*_) *−ln*(*K*_*M*_), one can conclude that s = ln(v) also follows approximately a distribution similar to **Eqs (8-9)** as *w*(*s, χ*) *≅ χexp*(*s*)(*χ+ exp*(*s*))^*−2*^with mean *s*_*m*_ *≅ ln*(*χ*) and variance *ϕ*_*s*_ *> ϕ*_*z*_ *> ϕ*_*y*_. Clearly, the mean values of (*γ, u, v*) depend strongly on (*κ, θ, χ*). Whereas, their variances are independent of them. **Eqs (7-9)** are the central results which describe the theoretical distributions of (*γ, u, v*) and their log-transforms (*y, z, s*) associated with K_M_, k_cat_ and K_E_ respectively.

### COMPUTATIONAL DATA ANALYSIS

To check the validity of **Eqs (7-9)**, we considered the BRENDA database values on K_M_, k_cat_, k_E_ and K_l_ of various enzymes across different substrates and experimental conditions [18-20]. Among these parameters, only K_M_ can be estimated directly from the experimental data using the double reciprocal plot or least square fitting procedures and other parameters should be estimated indirectly [3]. Particularly, k_cat_ will be estimated by dividing v_max_ = k_cat_ E_T_ by the total enzyme concentration (E_T_). K_l_ must be estimated from the data on the variation of K_M_ at different inhibitor concentrations. As a result, one can expect more external noise in the estimation of k_cat_, k_E_ and K_l_ than K_M_ along with inherent variations predicted by **Eqs (7-9)**. Further, the parameter values in this database with respect to a given enzyme are obtained under various experimental conditions that is essential to explore all the possible values of the random variables (*γ, u, v*) which otherwise will be generally restricted within a range by the natural selection pressure. To avoid such artefacts, unlike the analysis presented in Ref. [5], we considered all the values in the database irrespective of whether they are obtained using natural or synthetic substrates of enzymes. The latest BRENDA data file was downloaded from https://www.brenda-enzymes.org/download.php (version 2023_1). Histograms of the log transformed parameters of a given enzyme were constructed and the respective theoretical frequencies in a given bin range of the histogram were computed as follows.

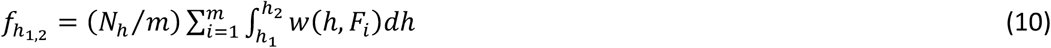

In this equation, the bin range of the variable *h* = (y, q, z, s) is (h_1_, h_2_) where *h*_l_ < *h*_2_ with the theoretical frequency 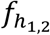, N_h_ is the total number of data points available for the random variable *h, H* is the control parameter (F = κ in case of y and q, F = θ in case of z and F = χ in case of s) and *w*(*h, F*) is the corresponding probability density function. Here m represents the number of density function components present in the sample. Sample variances were computed as 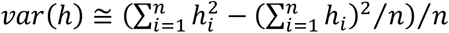 where n is the sample size. Multimodal type distributions will be observed when m > 1.

## RESULTS AND DISCUSSION

BRENDA database version 2023_1 has totally 7753 number of annotated enzyme entries out of which only 4990 entries have K_M_ values. Out of these, 10 entries have sample size of > 500 and 778 entries have sample size of > 50. We considered those entries with sample size > 50 for constructing the distribution of sample variances of y. We used those entries with sample size > 500 to check the validity of **Eq (8)**. Theoretical frequencies were constructed using **Eq (10)**. There were 2040 number of enzyme entries with K_l_ values out of which 189 entries were there with the sample size of > 50. We used these values to construct the distribution of sample variances of q. Processed datasets are available in the supporting materials.

Analysis results on ln(ϒ) values from BRENDA database and the theoretical predictions of **Eq (8)** are shown in **Fig 2**. Out of 10 enzyme entries with a sample size of > 500, glutathione transferase, beta-glucosidase, tyrosinase are shown in **Figs 2A, 2B** and **2C** respectively along with the prediction by **Eq (8)** with appropriate single fit value of κ. Enzymes such as laccase (EC: 1.10.3.2) and beta-lactamase (EC: 3.5.2.6) also exactly obey the prediction by **Eq (8)** with single κ i.e., m = 1 in **Eq (10)**. Close observation suggests that a normal density function of type 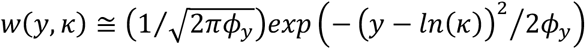 with *ϕ*_*y*_ *= π*^*2*^*/*3 also approximately fits these datasets. This means that a log-normal type density function can also be used to fit ϒ. However, there is no physical basis for the origin of such density function. Remarkably, alcohol dehydrogenase (EC: 1.1.1.1) exhibited a bimodal type density function (m = 2 in **Eq (10)**) which fits well with the average over two different κ values on **Eq (8)** as shown in **Fig 2D**. Enzymes such as carboxyl-esterase and ribulose bisphosphate carboxylase deviate from the prediction by **Eq (8)**. The distribution of sample variances of ln(ϒ) and ln(η) are shown in **Figs 2E** and **2F**. In line with **Eq (8)**, most probable value of the sample variance occurred around *ϕ*_y_ = π^2^/3 for y and around *ϕ*_q_ = π^2^ for q.

**FIGURE. 2.**
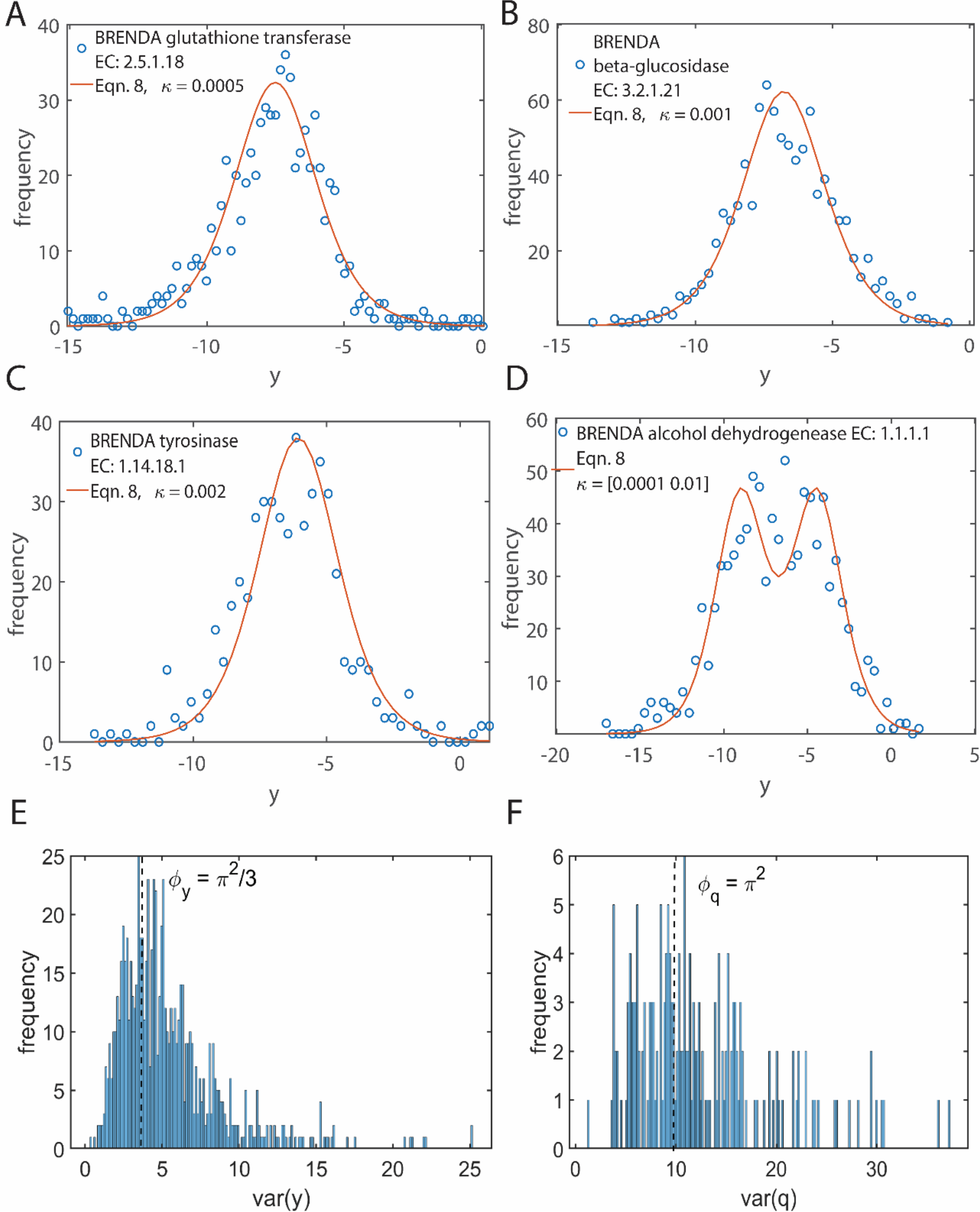
Comparison of the theoretical distribution of K_M_ (μ mol/lit) with the BRENDA database. **A-D**. Theoretical frequencies were computed using **Eq (8)** and **Eq (10)**. The default unit of K_M_ in BRENDA is mM and to convert it into M, K_M_ entries were multiplied by 10^-3^ and then log transformed as y = ln(ϒ) where K_M_ = < ϒ >. Total number of bins in histogram was set to 50. **A**. the total frequency for EC: 2.5.1.18 was 720 and bin size was 0.359. Here K = 0.0005.**B**. the total frequency for EC: 3.2.1.21 was 947 and bin size was 0.264. Here K = 0.001. **C**. the total frequency for EC: 2.14.18.1 was 504 and bin size was 0.31. **D**. the total frequency for this enzyme EC: 1.1.1.1 was 947 and bin size was 0.381. This enzyme exhibits a bimodal type density function with two distinct K values so that **Eqn. 10** was used with m = 2. **F-G**. Total number of bins was set to 200. **F**. histogram of var(y) with maximum around at ϕ _y_ = π ^2^/3. **G**. histogram of var(q) with maximum around at ϕ _q_ = π ^2^.

There were 2246 (189 with > 50 sample size, none with > 500 and 4 with > 250) number of enzyme entries with k_E_ values and 3370 (391 with > 50 sample size and 4 with > 500) enzyme entries with k_cat_ values. Observed distributions of z = ln(u) and s = ln(v) are shown in **Figs 3A-B** and **3C-D** respectively. The respective distributions of the sample variances (*ϕ*_*s*_, *ϕ*_*z*_) are shown in **Fig 3E-F**. Remarkably, the distributions of both z and s fit to bimodal type with m = 2 in **Eq 8** and **Eq (10)**. Such bimodal type of density function of z must be originating from the conformational fluctuations in the enzyme active site as observed in the single molecule experiments [21, 22]. The bimodal type density function of z = ln(u) will obviously reflect in the distribution of s = ln(v) since v = u / ϒ.

**FIGURE. 3.**
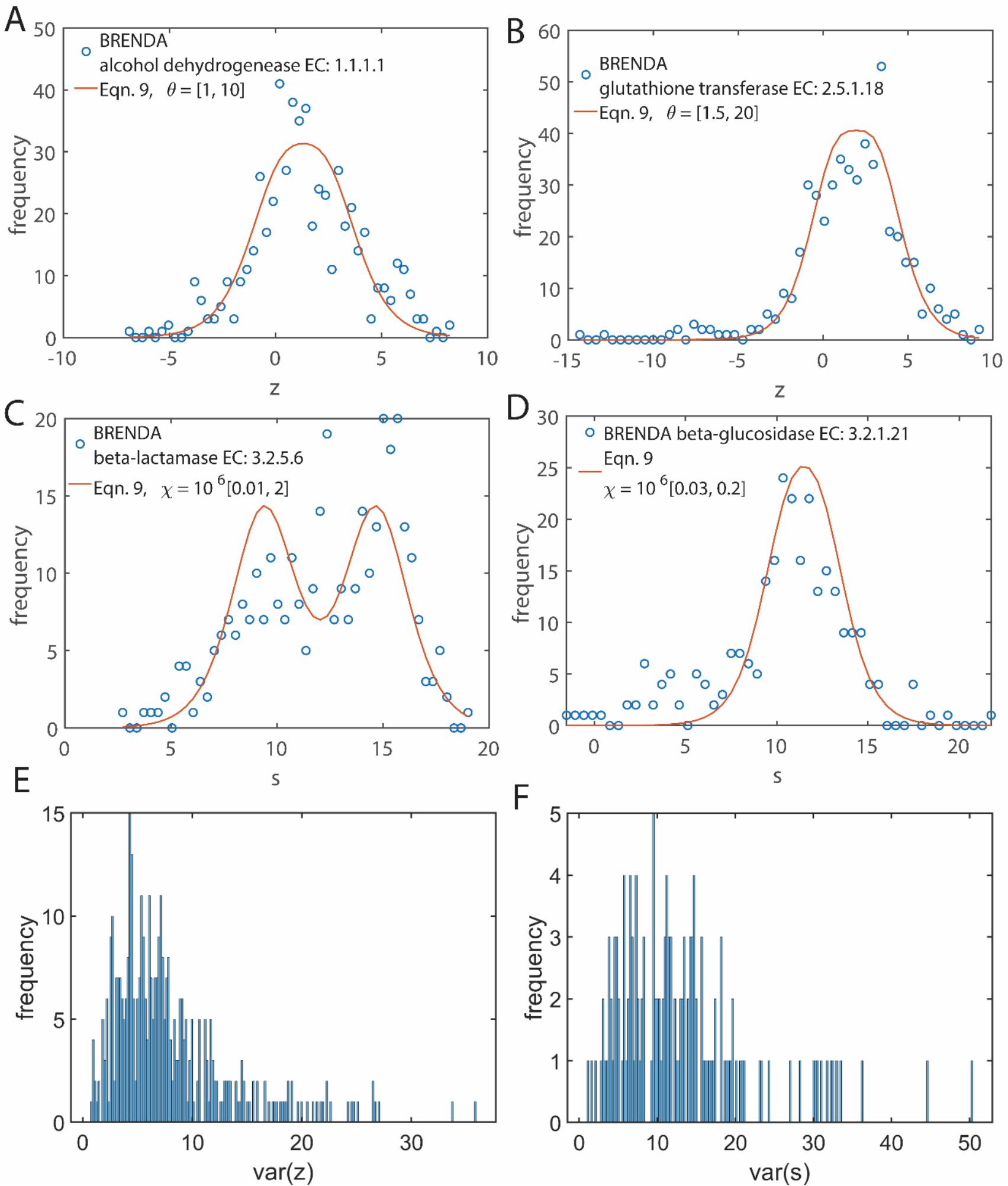
Comparison of the theoretical distribution of k_cat_ (1/second) and k_E_ (1/mol/lit/second) with the BRENDA database. **A-D**. Theoretical frequencies were computed using **Eq (9)** and **Eq (10)**. The default unit of k_E_ in BRENDA is 1/mM/lit/s and to convert it into M, k_E_ entries were multiplied by 10^3^ and then log transformed as s = ln(v) where k_E_ = <v>. Total number of bins was set to 50. **A**. the total frequency for EC: 1.1.1.1 was 559 and bin size was 0.3. Here m = 2 with 8 = [1, 10]. **B**. the total frequency for EC: 2.5.1.18 was 502 and bin size was 0.48. Here m = 2 with 8 = [1.5, 20]. **C**. the total frequency for EC: 3.5.2.6 was 340 and with bin size 0.332. Here m = 2 with x = 10^6^[0.1, 2]. **D**. the total frequency for EC: 3.2.1.21 was 264 and total number of bins was set to 50 with bin size 0.475. Here m = 2 with χ = 10^6^ [0.03, 0.2]. **F-G**. Total number of bins was set to 200. **F**. histogram of ϕ _2_. **G**. histogram of ϕ _s_.

The distribution of enzyme kinetic parameters is generally assumed to be dictated by the natural selection pressure [23, 24] which always works towards maximizing the substrate binding strength, product turnover and catalytic efficiency well within the framework of metabolism. There are two different aspects of these distributions viz. their curve geometry and the location of the peak. The shape of the density functions are mostly determined by the underlying biophysical aspects of the catalytic mechanism rather than the evolutionary pressure. This means that the parameters of these density functions (*κ, θ, χ*) which dictate the location of peaks are mainly influenced by the evolutionary selection pressure. Whereas, the curve geometry is determined by the underlying biophysical aspects of the catalytic mechanism. This is in fact supported by the observation that the variances of the enzyme kinetic parameters are independent on the parameters related to the respective density functions especially in the log-transformed space. The results presented here are essential to further understand how these enzyme kinetic parameters are designed and optimized by the natural selection process to work at maximum efficiency within the framework of metabolism.

## CONCLUSION

For the first time in the literature, we have derived the power-law type density functions of the enzyme kinetic parameters purely based on the biophysical arguments and successfully validated our findings with several examples on the available datasets. We have demonstrated that the experimentally observed enzyme kinetic parameters related to the substrate binding, product turnover and catalytic efficiency follow power-law type density functions. These finding are validated with the available BRENDA datasets on various enzymes across different substrates. The product turnover rates and catalytic efficiencies follow a bimodal type density functions in line with the single molecule experiments on enzyme catalysis which can be explained by a sum of two different power-law type density functions. Our results suggested that the curve geometry of the density functions is decided by the underlying biophysical factors and the location of the peak is dictated by the natural selection pressure.

## Notes

### Competing Interest Statement

The authors have declared no competing interest.

